# Neural Networks Estimate Muscle Force in Dynamic Conditions Better than Hill-type Muscle Models

**DOI:** 10.1101/2024.03.12.584454

**Authors:** Maria Eleni Athanasiadou, Monica A. Daley, Anne D. Koelewijn

## Abstract

Hill-type muscle models are widely used, even though they do not accurately represent certain muscle mechanics. We explored neural networks to develop new muscle models. We trained neural networks to estimate muscle force from activation, muscle length, and muscle velocity. Training data was recorded using sonomicrometry, electromyography, and a tendon buckle on two muscles of guinea fowl. First, we compared the neural network to a Hill-type muscle model, using the same data for network training and model optimization. Second, we trained neural networks on large datasets, in a more realistic machine learning scenario. We found that neural networks generally yielded higher coefficients of determination and lower errors than Hill-type muscle models. Our neural networks performed better when estimating forces on the muscle used for training, but on another bird, than on a different muscle of the same bird, which could be explained by inaccuracies in activation and force scaling. We extracted force-length and force-velocity relationships from the trained neural networks and found that both effects were underestimated and that both relationships were not replicated well outside of the training data distribution. We discuss suggested experimental designs to collect suitable training data and conclude that neural networks could provide an accurate alternative to Hill-type muscle models, particularly for modeling dynamic muscle behavior that is prevalent in faster movements, given a suitable training dataset, while scaling of the training data should be comparable between muscles and animals.

**Summary:** Neural networks predict muscle forces more accurately than Hill-type muscle models, particularly under dynamic conditions. However, they struggle to replicate the force-length and force-velocity relationships well.

## 1 Introduction

The human musculoskeletal system has sparked scientific interest for centuries, since knowledge of internal body forces is crucial in clinical assessments (Steele et al., 2012), prosthesis design (Ferris et al., 2012), and injury prevention (Matijevich et al., 2019), among others. However, direct measurements of these forces require highly invasive sensors and procedures, and are therefore generally not performed in humans, especially not during movement. Instead, scientists have focused on the musculoskeletal system of other species to understand the mechanics of muscles and other tissues. For example, A. V. Hill deciphered the working of muscles using frog specimens at maximum activation (Hill, 1938). Based on his research, the so-called Hill-type muscle models have been developed, which describe muscle mechanics and can be used to simulate muscle behavior. These Hill-type models are widely applied to answer research questions in humans (e.g., Steele et al. 2012; Koelewijn and Selinger 2022) and other animals (e.g., Hill 1938; Lee et al. 2013; Lemaire et al. 2016). They fall under the phenomenological model category, meaning that they mathematically describe an observed relationship, in this case between force development, muscle length, and muscle velocity, without necessarily describing the actual (chemical and mechanical) process that happens in the muscle. This relationship has since been expanded to include muscle activation (Hatze, 1977).

Originally, Hill-type muscle models consisted of two components representing the active muscle fibre and passive tissue in series (mainly the tendon) (Miller, 2018). The contractile element (CE) describes the behavior of active muscle fibre, mainly the force-length-velocity relationship (Zajac, 1989). This relationship is based on the sliding filament theory (Gordon et al., 1966). At a single fiber level, when a muscle is activated, crossbridges are formed between myosin and actin filaments. The formation of these crossbridges leads to the filaments sliding across one another, causing the muscle length to change and force to be generated. The amount of force that is being generated depends on the overlap of filaments and the rate of crossbridge cycling relative to the sliding speed. Besides the CE, these original models also commonly included a series elastic element (SEE), which represents the passive tissue connected in series to the active muscle tissue, mainly the tendon and aponeurosis and includes the compliance of fibers (Miller, 2018).

Since these original models, many different Hill-type muscle models have been formulated with different complexity levels. Usually, complexity was increased in order to incorporate more complex muscle behaviors and thereby increase accuracy. A common addition is a parallel elastic element (PEE), which represents the passive tissue parallel to the active muscle fibre (Miller, 2018). Another well-known extension was Zajac’s integration of muscle geometry and activation dynamics, which allowed for force prediction under various different physiological conditions (Zajac, 1989). Additional complexities like muscle fiber distribution (Herzog, 1999), histology (Lee et al., 2013), and orientation (Van den Bogert et al., 2011), as well as dynamic length-tension relationships (Herzog, 1999) have been incorporated in other more advanced Hill-type models. Besides increasing accuracy, the computational efficiency of Hill-type models in large-scale simulations has also been addressed by employing implicit numerical methods (Van den Bogert et al., 2011).

Despite the widespread use of Hill-type muscle models, it is known that these models do not fully capture muscle behavior as recorded experimentally. The isometric force-length and isotonic force-velocity relationships are central components of Hill-type models. These relationships were formulated under tightly controlled experimental conditions, where length and tension were kept constant, respectively. Therefore they rarely represent typical in-vivo conditions, leaving room for uncertainty as to whether, or to what extent, they accurately reflect dynamic in-vivo muscle behavior. Another important aspect of muscle mechanics that the Hill-type model does not capture is the tension change in muscle following a stretch or a shortening (Abbott and Aubert, 1952). It has been suggested that titin, a third filament, winds around the thin, actin filament during crossbridge formation, and thereby acts as a spring, creating force enhancement during active stretch (Nishikawa et al., 2012). During active shortening, the titin filament unwinds, thereby causing force depression during shortening (Nishikawa et al., 2012). Other aspects that are not commonly included in the Hill-type muscle model are changes in optimal fibre length and maximum shortening velocity with activation level. Roszek and Huijing and Rack and Westbury have shown that the optimal fibre length changes with the muscle activation levels (Roszek and Huijing, 1997), while Chow and Darling showed the same for the maximum shortening velocity (Chow and Darling, 1999). These Hill-type muscle model limitations could be related to the fact that the model is applied to different activation levels, while it is based on experiments performed at maximum activation (Hill, 1938).

To overcome these limitations, in recent years, further extensions have been proposed to the Hill-type muscle model (e.g. Chow and Darling 1999; McGowan et al. 2013), while completely new models have also been proposed (e.g., Tahir et al. 2018; Millard et al. 2023). For example, the history-dependent Hill-type model (McGowan et al., 2013) incorporates the force enhancement or depression caused by the muscle being stretched or shortened (Miller, 2018; Abbott and Aubert, 1952), respectively. Another extension includes the relationship between maximum shortening velocity and muscle stimulation (Chow and Darling, 1999; Nitschke et al., 2020). Instead of extending the Hill-model, Whitney et al. (2022) and Millard et al. (2023) have developed completely new phenomenological models to improve muscle model accuracy. These models are based on the mechanics of the three muscle filaments: actin, myosin, and titin (Nishikawa et al., 2012; Millard et al., 2023), while Hill-type models only include the actin and myosin filaments. Both models (Millard et al., 2023; Whitney et al., 2022) outperform the Hill-type model, yet their force estimations could still be improved further (Millard et al., 2023), and setting their parameters can be difficult (Whitney et al., 2022).

Instead of further improving these existing phenomenological models, an alternative approach is to use machine learning to develop a model completely based on in-vivo muscle mechanics data, gathered during different activities (Liu et al., 1999). In theory, a large and rich enough dataset containing such data should enable us to train a neural network that replicates muscle mechanics relevant to movement and is more accurate than a model based on ex-vivo experiments. In practice, obtaining such a dataset on humans is virtually impossible, since sensors should be implanted into the muscles to record muscle length, force, and activation. However, such datasets have been collected for guinea fowl (Daley and Biewener, 2011) and goats (Lee et al., 2013), among others. Similar to a Hill-type model, which is normalized to optimal fibre length and maximum isometric force to be applicable to different muscles and species, a neural network trained on such a dataset might also be applicable to other muscles and species, as long as it is appropriately normalized. However, to do so, it should be validated that the trained network correctly represents muscle mechanics. Previously, an artificial neural network was successfully trained to estimate muscle force from electromyography data of a cat soleus (Liu et al., 1999). However, the relationship between electromyography and muscle force is not always direct, since the muscle length and velocity also affect force generation (Nishikawa et al., 2007). This relationship is especially well visible in a dataset containing perturbations, such as in the dataset by Daley and Biewener (2011).

Therefore, in this paper, we developed neural networks to estimate muscle force from muscle length, velocity, and activation. We trained those neural networks on existing data from guinea fowl (Daley and Biewener, 2011). First, we trained a neural network on a smaller dataset and compared its accuracy to that of a state-of-the-art Hill-type muscle model, which was optimized using the same data that was used to train the neural network. For this comparison, we chose a two element Hill-type muscle model with only a forcelength-velocity relationship, rather than a more complex model. Such complex models are commonly used to investigate muscle mechanics of individual muscles (Zahalak, 1986; McGowan et al., 2013), but they are not necessarily more effective than Hill-type muscle models (Lemaire et al., 2016). Second, we investigated the accuracy of networks that were trained on larger datasets and thus a more realistic machine learning scenario. We first evaluated the networks by comparing their estimated muscle forces to the measured ones and then explored if the networks replicated known muscle mechanics, specifically the force-length and force-velocity relationships. We investigated if these relationships emerge from the networks, which would mean that they correctly represent muscle mechanics. Therefore, this comparison could provide an additional validation of the neural networks’ ability to fully capture muscle behavior, which is necessary to confidently use these models across species.

## 2 Materials and Methods

We used an existing dataset that was recorded on guinea fowl (Athanasiadou et al., 2025), containing activation, muscle length, and muscle velocity as inputs and muscle force as output. We used this dataset for network training and Hill-type muscle model optimization, and evaluated how well neural networks and Hill-type muscle models could estimate forces on the same muscle and bird as used for training, a different muscle of the same bird, the same muscle of a different bird, and a different muscle of a different bird. All code used for neural network training, Hill-type muscle model optimization and analysis can be found here (Athanasiadou et al., 2024).

### 2.1 Guinea Fowl Data

The dataset contained data from six guinea fowl. We summarize the protocol here and refer readers to Daley and Biewener (2011) for a more detailed description. Briefly, this dataset contained between 12 and 15 walking and running trials of about 15 to 30 seconds for each bird. The measurements were taken during level and obstacle trials between 1.8 to 4.5 m s^-1^. During the obstacle trials, the birds would suddenly step on a raised obstacle (5 or 7 cm) every four to five steps. Data was recorded using a tendon buckle (force), sonomicrometry (contractile element length), and electromyography (activation) for two different muscles; the lateral head of the gastrocnemius (LG) and the digital flexor to the lateral toe (DF). The contractile element velocity in the dataset was calculated as the time derivative of the contractile element length. Out of the six guinea fowl, we used the data of both muscles for three birds. For the fourth bird, the data quality of the LG was not very good due to cross-talk noise in the electromyography and the quality of the sonomicrometry for the DF was poor on day 1. Therefore, we did not include the DF data recorded on day 1 and the LG data in its entirety for bird 4. We only used the LG data for the fifth bird, since the electromyography failed for the DF. We omitted the sixth bird entirely due to data quality issues. In previous studies (e.g. Daley and Biewener 2011), a sufficient sample of individual strides with good data quality were identified for each bird, but this approach is not suitable here, because training neural networks requires as much data as possible and thus entire trials and consecutive strides rather than individually picked gait cycles.

We used the tendon force in N, the normalized contractile element length and velocity, and the raw electromyography signal and processed the data as follows. First, we computed the maximum isometric force of each bird’s LG and DF using the physiological cross-sectional area (Daley and Biewener, 2011), and assuming a maximum stress of 0.36 N mm^-2^ (Cox et al., 2019), which is within the commonly accepted range of 0.26 to 0.4 (Ker et al., 1988; Wells, 1965; Jayes and Alexander, 1982; Head et al., 2011). We then normalized the measured tendon force using this maximum isometric force. Then, we processed the electromyography data similar to Whitney et al. (2022): we first applied a third order Butterworth high-pass zero-phase digital filter with a cut-off frequency of 30 Hz, then we rectified the signal, after which we applied a third order low-pass zero-phase digital filter with a cut-off frequency of 6 Hz. This activation signal was then normalized to the maximum measured activation in the processed data over all trials of the specific bird and muscle, and a time delay of 23.6 ms was added to account for excitation contraction coupling (Daley and Biewener, 2011). The same 6 Hz low-pass filter was also applied to the contractile element velocity.

### 2.2 Neural Network Training

We designed several neural networks to estimate tendon force from activation, contractile element length, and contractile element velocity. To replicate a Hill-type muscle model as close as possible, without introducing any bias in favor of the neural networks (e.g. by incorporating history dependence), these networks were set up to calculate the tendon force at each time point individually from the inputs at the same time point. First, we compared two neural networks to the Hill-type muscle model and trained them on the same data that was also used to optimize the Hill-type muscle model parameters. Due to the computational cost related to the model optimization, we made this comparison on a smaller dataset, consisting of a single trial. Specifically, we trained a network using the data of the LG of the first trial of bird 1, a level trial at 1.8 m s^-1^. The second network was trained on the twelfth trial of the same bird and muscle, a trial with 7 cm obstacles at 4.5 m s^-1^. Second, we trained neural networks on larger datasets to investigate their prediction accuracy in a more commonly used machine learning scenario. We used a leave-one-out training approach and trained four networks, excluding the LG data of a different bird for each, so that each network was trained on the LG data of three birds. We call these networks NN-b1 (bird 1 was left out), NN-b2 (bird 2 was left out), NN-b3 (bird 3 was left out) and NN-b5 (bird 5 was left out). The number of trials used for training ranged from 39 to 41.

We used the following process to train the networks. First, we split the data into three categories; the *training* data, on which the neural network was trained, the *validation* data, to validate the hyperparameter optimization, and the *test* data, on which the trained neural network made predictions. The training data contained the first 80% of all trials used for training, while the validation was performed on the last 20%. This split was chosen to ensure that the training data did not include information from a later time point than any of the validation data, which could lead to data leakage (Dorschky et al., 2023). We normalized the complete dataset to have a mean of zero and a standard deviation of one. We optimized the network’s hyperparameters, specifically the number of layers, number of nodes in each layer, and the activation function of the hidden layers, using the automatic hyperparameter optimization that is included in the MATLAB function fitrnet, which uses a Bayesian optimization. We repeated this hyperparameter optimization 10 times when training on a single trial, and five times when training on the large dataset, to account for random effects of the Bayesian optimization, and chose the network with the smallest loss overall. We performed all training and analysis in MATLAB R2022a and R2022b (Mathworks, Natick, MA, USA).

### 2.3 Hill-type Muscle Model and Optimization

We used the Hill-type muscle model described by Zajac (1989), Van den Bogert et al. (2011), and Herzog (1999). We chose this model, since its mathematical equations are such that the shape of the force-length and force-velocity relationship is preserved during an optimization. We found that for other models, the equations could be optimized such that the shape of the force-length and force-velocity relationship changed. We determined the tendon force, *F*_*Hill*_, as the sum of the force in the contractile element, *F*_*CE*_, and the parallel elastic element, *F*_*P EE*_, using the following relationships:

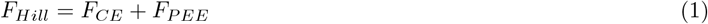

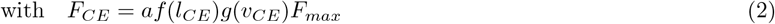

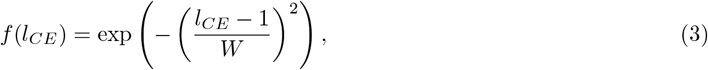

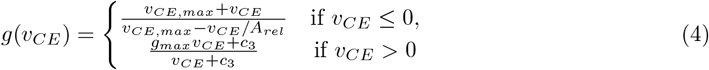

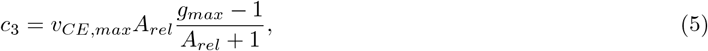

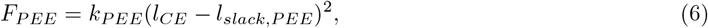

where *f* (*l*_*CE*_) is the force-length relationship with width, *W, g*(*v*_*CE*_) the force-velocity relationship with maximum shortening velocity, *v*_*CE,max*_, the curvature of the force-velocity relationship, *A*_*rel*_, the maximum force amplification during lengthening, *g*_*max*_, and factor *c*_3_, which was calculated such that the force-velocity relationship is continuous, *k*_*P EE*_ the stiffness of the parallel elastic element, and *l*_*slack,P EE*_ its slack length. Note that the muscle length, *l*_*CE*_, and velocity, *v*_*CE*_, are normalized to optimal fibre length.

We optimized the Hill-type muscle model using the same data as used for network training, to allow for a fair comparison between the model and the neural network. We optimized the width of the force-length relationship, *W*, the maximum shortening velocity, *v*_*CE,max*_, the curvature of the force-velocity relationship, *A*_*rel*_, the maximum force amplification during lengthening, *g*_*max*_, the stiffness of the parallel elastic element, *k*_*P EE*_, and its slack length, *l*_*slack,P EE*_. As an objective function, we used the mean square difference between the calculated, *F*_*Hill*_, and measured, *F*_*meas*_, muscle force:

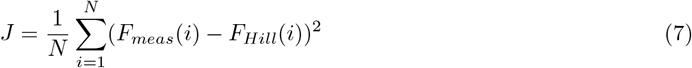

We optimized the parameters using covariance matrix adaptation, evolutionary strategy (CMA-ES) (Hansen and Ostermeier, 2001), using 50 iterations. We performed the optimizations in MATLAB R2021a (Mathworks, Natick, MA, USA).

### 2.4 Data Analysis

First, we investigated if the neural network was able to outperform the Hill-type muscle model. To do so, we optimized the Hill-type muscle model on the same data of bird 1 as was used for neural network training. We trained two neural networks and optimized the Hill-type muscle model twice: once using the first trial at 1.8 m s^-1^ and a level surface and once using the twelfth trial at 4.5 m s^-1^ with 7 cm obstacles. We call these models NN-r01, NN-r12, Hill-r01, and Hill-r12, respectively. We compared all models using the root-mean-square error (RMSE), normalized to maximum isometric force, and the coefficient of determination (R^2^) between the measured and estimated (either through the neural network or Hill-type muscle model) force. We compared both metrics on the same bird and muscle as used for training, on all trials that were not used in training, on the same bird’s DF, to investigate both models’ ability to predict force of another muscle, on bird 2’s LG, to investigate the models’ ability to predict force of another bird, and on bird 2’s DF, to investigate the prediction accuracy when both bird and muscle were different.

Second, we investigated the accuracy of neural networks in a more common training approach using much larger datasets. For the birds that were included in each training, we used the recordings of the LG of all but one trial of each bird, which was excluded for testing. All trials of the LG muscle of the bird that was left out of training were also used for testing the network. All networks were compared using the RMSE, the relative RMSE (RMSE%) and the coefficient of determination (R^2^) between the measured force and the networks’ predictions. The relative RMSE was calculated using the maximum measured force during each trial. The testing approach was similar as before, evaluating each network’s estimation accuracy on four scenarios: on the same bird and muscle as used in training, on the same bird but another muscle (DF), on the same muscle of a different bird, and on another muscle (DF) of a different bird. Bird 4 was never used when training the networks, since its LG recordings were of low quality. Therefore, predictions for its DF muscle were always made based on a network that was trained on other birds. Similarly, no predictions were made for the DF muscle of bird 5, because no electromyography data was available.

Third, to investigate if the above networks replicated muscle mechanics, we compared the force-length and force-velocity relationships that emerged from the neural networks trained on a large amount of data to those of the Hill-type muscle model. To do so, we varied the fiber length between 60% and 140% of the optimal fibre length and calculated the resulting force normalized to maximum isometric force, assuming isometric conditions. The force-length relationship includes the PEE, since the CE and the PEE cannot be separated in a neural network. Then, we varied the contractile element velocity between -10 and 10 fibre lengths per second, assuming optimal fibre length, and again calculated the resulting force normalized to maximum isometric force. We repeated these for five different activation levels: 0.2, 0.4, 0.6, 0.8 and 1.

## 3 Results

### 3.1 Comparison of Neural Network to Hill-type Muscle Model

We found that the training and validation loss for NN-r01 were slightly lower than for NN-r12, while for both models, the training and validation loss show a similar trend (Fig. 1 and 2). After training, the neural network NN-r01 had a training loss of 0.055 and a validation loss of 0.13 after 29 iterations, while NN-r12 had a training loss of 0.16 and a validation loss of 0.20 after 44 iterations. Both models consisted of three hidden layers with rectified linear activation units, NN-r01 with 300, 254, and 200 nodes, and NN-r12 with 188, 140, and 13 nodes. We also found that similar results were achieved with vastly different network architectures, as the best networks found during hyperparameter optimization had similar losses but different characteristic features. For example, for NN-r12, the number of layers varied between two and three, the first layer ranged from 21 to 291 nodes, the second layer ranged between 10 and 297 nodes and the third layer between 13 and 245 nodes. Figures 1 and 2 also show some evidence of overfitting to the training data, as the training loss is lower than the validation loss.

**Figure 1:**
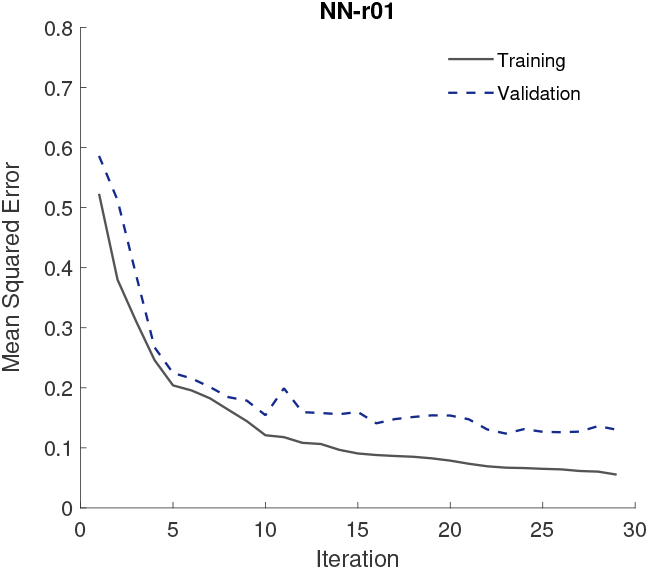
Training and validation loss during training of neural network NN-r01 (level at 1.8 m s^-1^).

**Figure 2:**
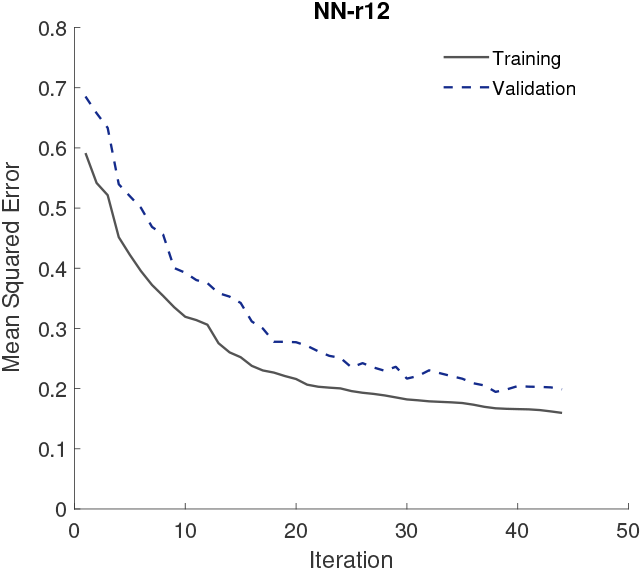
Training and validation loss during training of neural network NN-r12 (7 cm incline at 4.5 m s^-1^).

Our neural networks yielded lower mean RMSE than the Hill-type muscle models for all four test cases: when applied to the bird and muscle that were used for training and optimization, when applied to a different muscle, to a different bird, and when both are different (Table 1). Both neural networks produced lower mean RMSE values than the Hill-type muscle models, with the error reduction between 20%-30% for the LG and 30%-50% for the DF. The neural network NN-r12 had the lowest mean RMSE between the two neural networks on all four test cases. For the Hill-type muscle models, the mean RMSE of Hill-r01 was lower, except when the models were tested on the same muscle of a different bird (bird 2’s LG, Table 1).

**Table 1:**
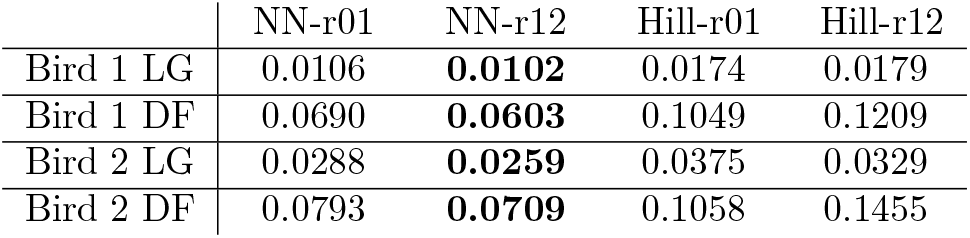
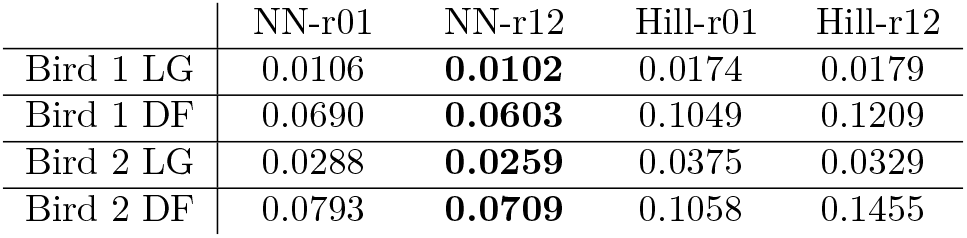
Mean root-mean-square-error (RMSE), normalized to maximum isometric force, between measured and estimated forces, averaged over all trials in the testing data. Bird 1 LG represents the same bird and muscle (lateral gastrocnemius, LG) as used for training, bird 1 DF the same bird, but a different muscle (digital flexor, DF), bird 2 LG a different bird, but same muscle, and bird 2 DF a different bird and a different muscle. Bold font indicates lowest mean RMSE.

|  | NN-r01 | NN-r12 | Hill-r01 | Hill-r12 |
| --- | --- | --- | --- | --- |
| Bird 1 LG | 0.0106 | **0.0102** | 0.0174 | 0.0179 |
| Bird 1 DF | 0.0690 | **0.0603** | 0.1049 | 0.1209 |
| Bird 2 LG | 0.0288 | **0.0259** | 0.0375 | 0.0329 |
| Bird 2 DF | 0.0793 | **0.0709** | 0.1058 | 0.1455 |

Similar to the superior performance in mean RMSE, the mean coefficient of determination of our neural networks was also higher than those of the Hill-type muscle models (Table 2). In all test cases, the neural network NN-r12 performed best, while the Hill-type muscle model Hill-r12 performed worst, except for the test on the same muscle of a different bird, where the model Hill-r01 performed worst. The mean coefficient of determination was negative for both Hill-type muscle models, for all test cases involving a different muscle. We found that the neural network estimation matched the measurements for all test cases, but especially well on the same bird and same muscle, indicating that there could be some overfitting to the training data (Fig. 3). Especially in Fig. 3(A) the neural network’s force estimates matched the measured data very well. The peak force in the DF muscle is underestimated by the neural network, while the Hill-type muscle model commonly overestimated the force in this muscle by approximately three times the mean value of the measured force (Fig. 3(B) and (D)). Such a significant overestimation or even underestimation of force, in which prediction errors exceed the variability of measured values, would explain negative values in the coefficient of determination. At the same time, the duration of the individual force peaks is longer in the neural network estimations than in the measurements. Using the LG muscle of a different bird, the Hill-type muscle model estimations were less smooth than the measurements and the neural network estimations, but the peaks’ magnitudes matched better, and so did the timing of force generation (Fig. 3(C)).

**Table 2:**
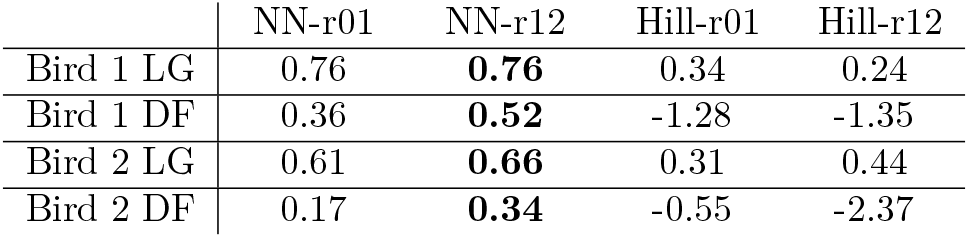
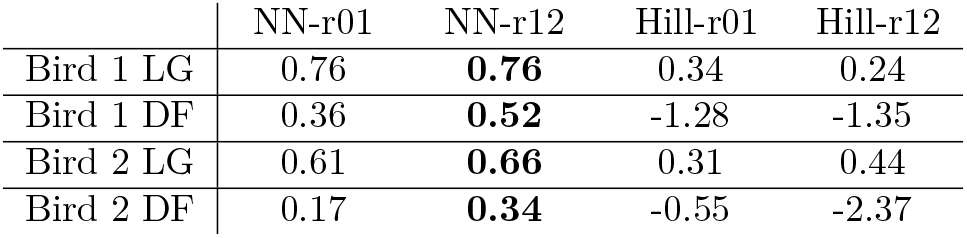
Mean coefficient of determination (R^2^) between measured and estimated forces, averaged over all trials in the testing data, for the same four test cases: the same bird and muscle as for training, the same bird but a different muscle than used for training, a different bird but the same muscle as used for training, and a different bird and different muscle. Bold font indicates highest mean coefficient of determination.

|  | NN-r01 | NN-r12 | Hill-r01 | Hill-r12 |
| --- | --- | --- | --- | --- |
| Bird 1 LG | 0.76 | **0.76** | 0.34 | 0.24 |
| Bird 1 DF | 0.36 | **0.52** | -1.28 | -1.35 |
| Bird 2 LG | 0.61 | **0.66** | 0.31 | 0.44 |
| Bird 2 DF | 0.17 | **0.34** | -0.55 | -2.37 |

**Figure 3:**
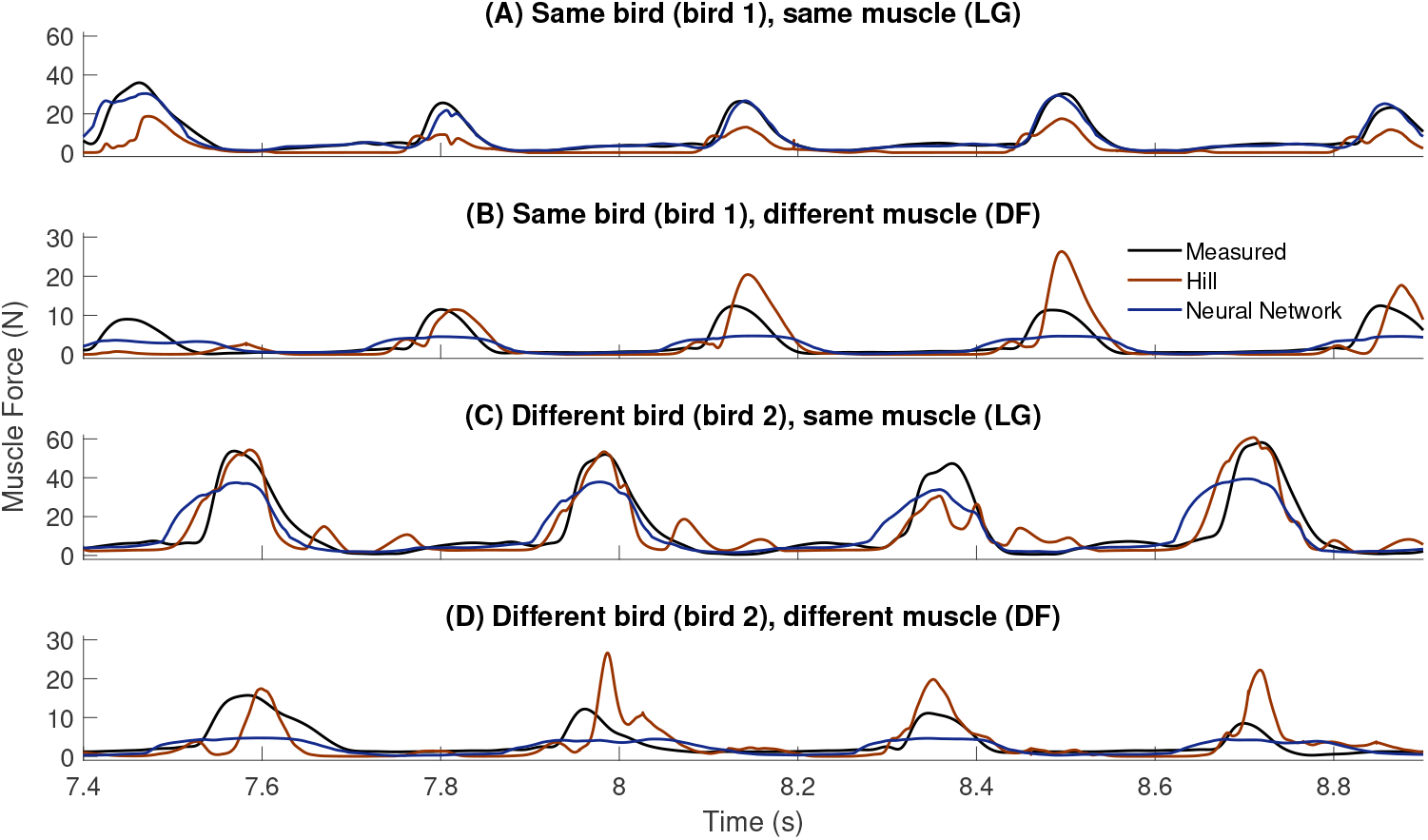
Muscle force over time for part of a trial with speed 3.8 m s^-1^ and 7 cm elevation for (A) the same bird and muscle as used for network training and Hill-type muscle model optimization, (B) a different muscle of the same bird, (C) the same muscle of a different bird, and (D) for a different muscle of a different bird. We plotted the results for the neural networks and Hill-type muscle models with the lowest mean RMSE, i.e., NN-r12 in (A)-(D), Hill-r01 in (A), (B), and (D), and Hill-r12 in (C).

### 3.2 Predictions with Neural Networks Trained on Large Datasets

The neural networks trained on large datasets combining different birds, had training losses that ranged from 0.13 to 0.25 and validation losses that ranged from 0.14 to 0.24 after 61 to 88 iterations. The validation losses of these networks were close to the losses of NN-r01 and NN-r12, while the training losses were higher than that of NN-r01 and closer to that of NN-r12. All of the networks trained on large datasets consisted of two layers with rectified linear activation units, except for the network NN-b1, which had three layers with hyperbolic tangent activation units. The networks’ architectures varied, with the first layer ranging from 81 to 300 nodes, the second layer 12 to 232 nodes and the third layer, of NN-b1, having 7 nodes. The difference between the training and the validation losses of each network trained on a large dataset, was smaller compared to the losses of the networks trained on fewer data. This indicates less overfitting to the training data, while the difference between training and validation loss again remained similar throughout the training of all networks, as Fig. 4 indicates for NN-b1.

**Figure 4:**
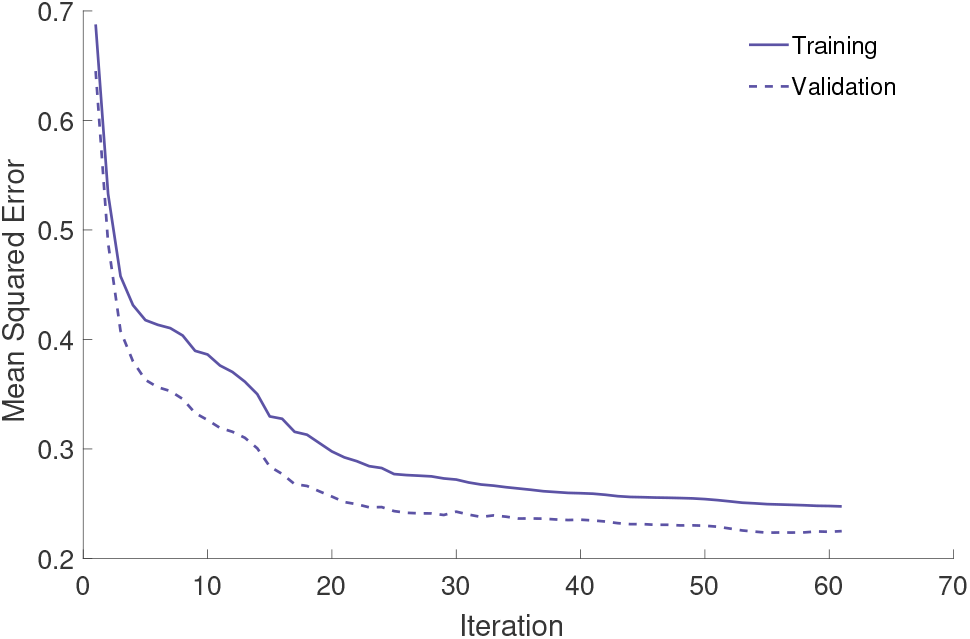
Training and validation loss during training of the neural network NN-b1.

Our networks generally predicted muscle force more accurately when predicting on the same muscle of a different bird compared to predicting on a different muscle of the same or a different bird, since the mean RMSE and the mean relative RMSE are higher, and the mean coefficients of determination are lower for the DF than for the LG (Table 3). NN-b5 is the exception, since the mean relative RMSE for the LG of bird 5 is higher than for the DF of all birds, while the mean coefficient of determination is negative and lower than that of the DF of all birds. Overall, all networks’ predictions are best for bird 1 for both the LG and DF. Even when bird 1 is left out of the training data, the mean RMSE of the LG is lowest and all three metrics for the DF are best for bird 1. The mean RMSE and mean relative RMSE are lowest or second lowest for bird 1 for both muscles, while the same goes for the mean coefficient of determination, except for the LG using NN-b5, where the mean coefficient of determination is the lowest for the LG of bird 1 among all networks. Conversely, when looking at the LG, which was used in training, for birds 2 and 5, the mean RMSE and mean relative RMSE are generally highest, even when these birds were part of the training dataset (NN-b1 and NN-b3). The only exception is the mean RMSE of bird 5 using NN-b3, which is smaller than that of bird 3. Comparing the performance within each muscle, we found that the magnitude of the mean RMSE for the muscle used for training (LG) was similar between the networks (between 0.01 and 0.02). However, the mean RMSE for the unseen muscle (DF) was at least twice as high for birds 3 and 4 compared to birds 1 and 2, though the difference was smaller for the mean relative RMSE, and even more pronounced for the mean coefficients of determination, where the coefficients were either negative or very close to zero for birds 3 and 4. Comparing the performance between the two muscles, the mean RMSE for the DF was overall at least twice as high as those of the LG, which was used for training. The magnitude of the mean coefficients of determination for the DF were generally below 0.5, with almost half of them being negative and only the mean coefficient for bird 1 using NN-b5 being 0.59. The mean coefficients of determination of the LG were generally above 0.5, except for bird 5, for which the highest mean coefficient of determination was 0.55 in NN-b2.

**Table 3:**
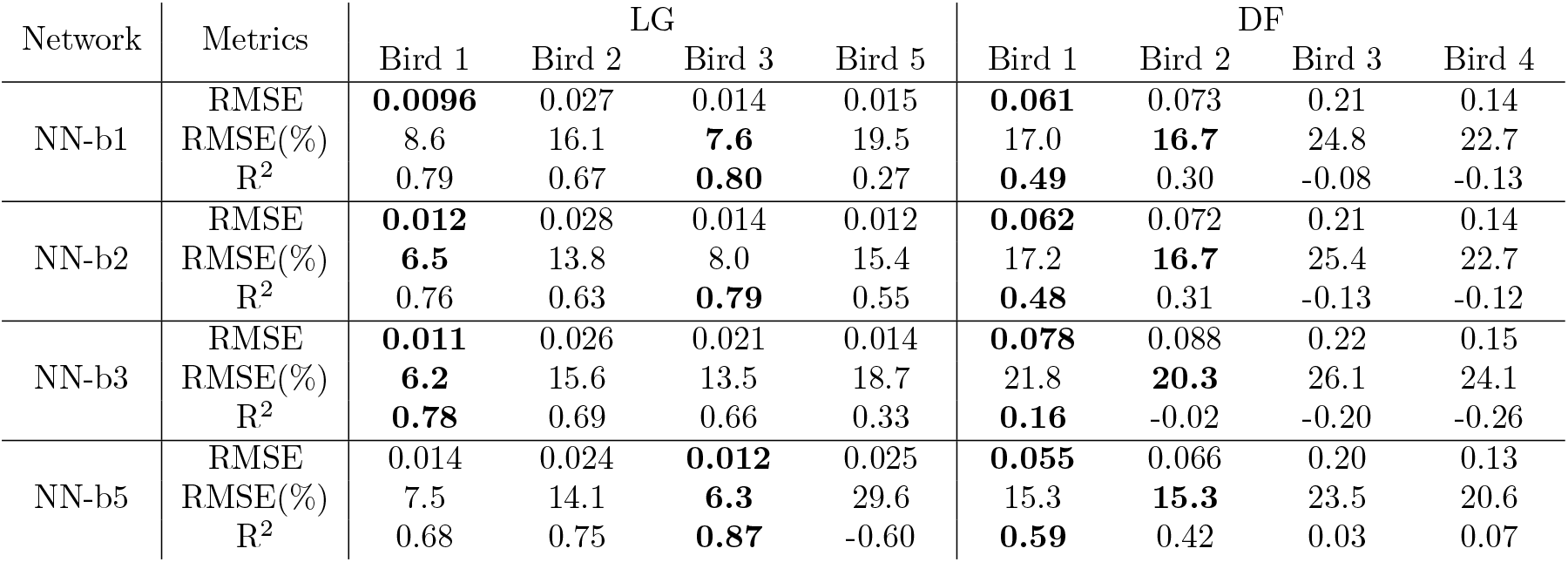
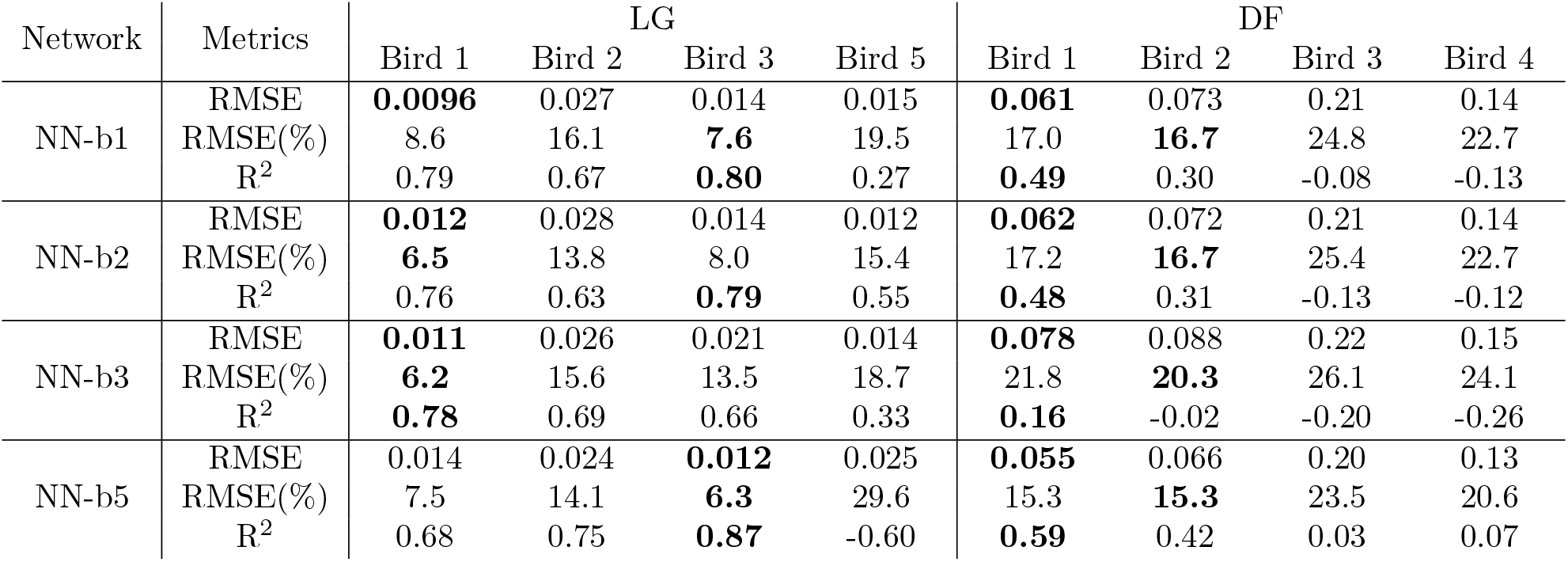
Mean RMSE, normalized to maximum isometric force, mean RMSE%, relative to maximum force, and mean coefficient of determination (R^2^) between force estimations and measurements, averaged over all trials in the testing data, for the neural networks trained on the LG data of different combinations of birds.

| Network | Metrics | LG |  |  |  | DF |  |  |  |
| --- | --- | --- | --- | --- | --- | --- | --- | --- | --- |
|  |  | Bird 1 | Bird 2 | Bird 3 | Bird 5 | Bird 1 | Bird 2 | Bird 3 | Bird 4 |
| NN-b1 | RMSE | **0.0096** | 0.027 | 0.014 | 0.015 | **0.061** | 0.073 | 0.21 | 0.14 |
|  | RMSE(%) | 8.6 | 16.1 | **7.6** | 19.5 | 17.0 | **16.7** | 24.8 | 22.7 |
| | $R^2$ | 0.79 | 0.67 | **0.80** | 0.27 | **0.49** | 0.30 | -0.08 | -0.13 |
| NN-b2 | RMSE | **0.012** | 0.028 | 0.014 | 0.012 | **0.062** | 0.072 | 0.21 | 0.14 |
|  | RMSE(%) | **6.5** | 13.8 | 8.0 | 15.4 | 17.2 | **16.7** | 25.4 | 22.7 |
| | $R^2$ | 0.76 | 0.63 | **0.79** | 0.55 | **0.48** | 0.31 | -0.13 | -0.12 |
| NN-b3 | RMSE | **0.011** | 0.026 | 0.021 | 0.014 | **0.078** | 0.088 | 0.22 | 0.15 |
|  | RMSE(%) | **6.2** | 15.6 | 13.5 | 18.7 | 21.8 | **20.3** | 26.1 | 24.1 |
| | $R^2$ | **0.78** | 0.69 | 0.66 | 0.33 | **0.16** | -0.02 | -0.20 | -0.26 |
| NN-b5 | RMSE | 0.014 | 0.024 | **0.012** | 0.025 | **0.055** | 0.066 | 0.20 | 0.13 |
|  | RMSE(%) | 7.5 | 14.1 | **6.3** | 29.6 | 15.3 | **15.3** | 23.5 | 20.6 |
| | $R^2$ | 0.68 | 0.75 | **0.87** | -0.60 | **0.59** | 0.42 | 0.03 | 0.07 |

Our neural networks trained on large datasets were again able to replicate the measured force well when applied to a muscle they were trained on, while for the other muscle, the peak force was underestimated (Fig. 5 for network NN-b1, Supporting Fig. **??, ??**, and **??** for other networks). In the cases where the peak force was severely underestimated (Figure 5(B), Supporting Fig. **??**(B) and **??**(B) & **??**(D)) or overestimated (Supporting Fig. **??**(C)), the mean coefficient of determination became negative. Figure 5(A) shows a trial that was excluded from training, for a bird and muscle that were used for training. Here the neural network estimations matched the measurements closely. The force was consistently underestimated slightly, while the small initial peaks (e.g. at 10.35 s and 10.95 s) were estimated more smoothly than in the data. For the predictions on the LG muscle of a different bird, the peak forces were over- or underestimated, while the smaller peaks were overestimated (10.7 s and 11.15 s) or of similar size (10.3 s and 11.6 s) (Fig. 5(C)). For the DF, which was not used in training, the neural network underestimated the peak force by at least 6 N for the unseen bird (bird 1) (Fig. 5(D)) and more than 20 N for bird 3 (Fig. 5(B)). Furthermore, the timing of the force does not match, with force generation having a shorter period for bird 3 (Fig. 5(B)) and a longer period for bird 1 (Fig. 5(D)).

**Figure 5:**
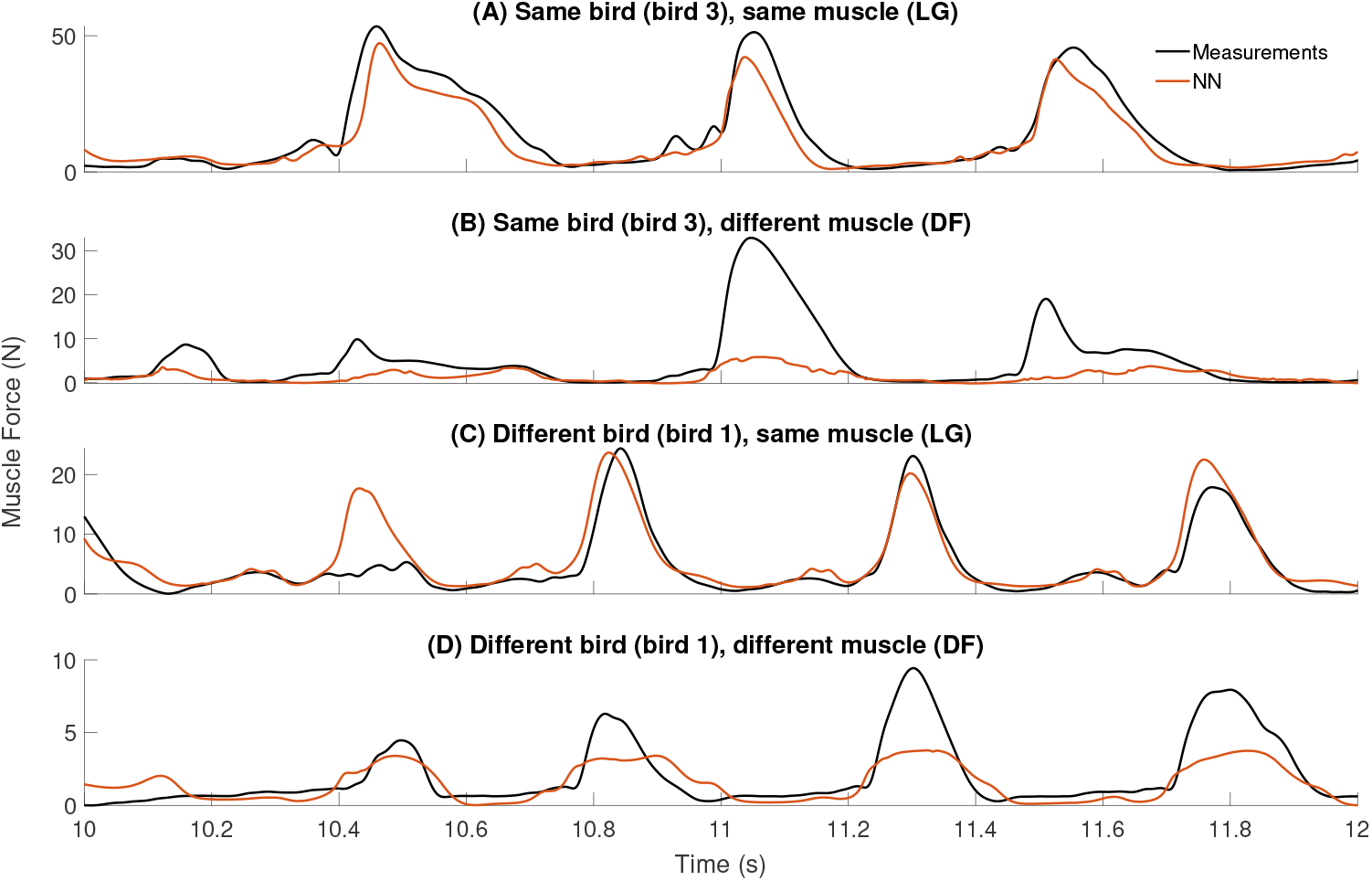
Muscle force predictions by NN-b1 and measurements over time, for part of a trial with speed 1.8 m s^-1^ and 7 cm elevation for (A) the same bird and muscle as used for network training, (B) the same bird but a different muscle, (C) the same muscle of a different bird, and (D) a different muscle of a different bird.

A visualization of this neural network’s force-length and force-velocity relationships can explain the underestimation of the DF force, and shows that these relationships were not represented well by the network (Fig. 6 for NN-b1. Supporting Fig. **??, ??**, and **??** for other networks). The maximum normalized muscle force was about 0.14 for both relationships, which indicates that the maximum muscle force is never achieved. This result indicates that the normalization to maximum measured activation in the processed data might not have been equivalent between the DF and LG, which explains the underestimations seen in Fig. 3 and Fig. 5. At 0.2 activation, the force-length relationship has a characteristic shape with a maximum at close to optimal fibre length, but with increasing activation levels, the maximum shifts to increasingly longer fibre lengths. At maximum activation, the descending slope is not visible anymore. The force-velocity relationship shows a decrease in muscle force generation with increasing shortening velocities, but the increased force generation during lengthening is not achieved. These force-length and force-velocity patterns are followed by NN-b3 and NN-b5 as well. For NN-b2, the force-length relationship maintains the maximum force close to the optimal fibre length for all activation levels, and a constant force-length relationship is observed at maximum activation. The force-velocity relationship better resembles the expected force-length relationship graph than the force-length relationship of NN-b1 does (Supporting Fig. **??, ??**, and **??**).

**Figure 6:**
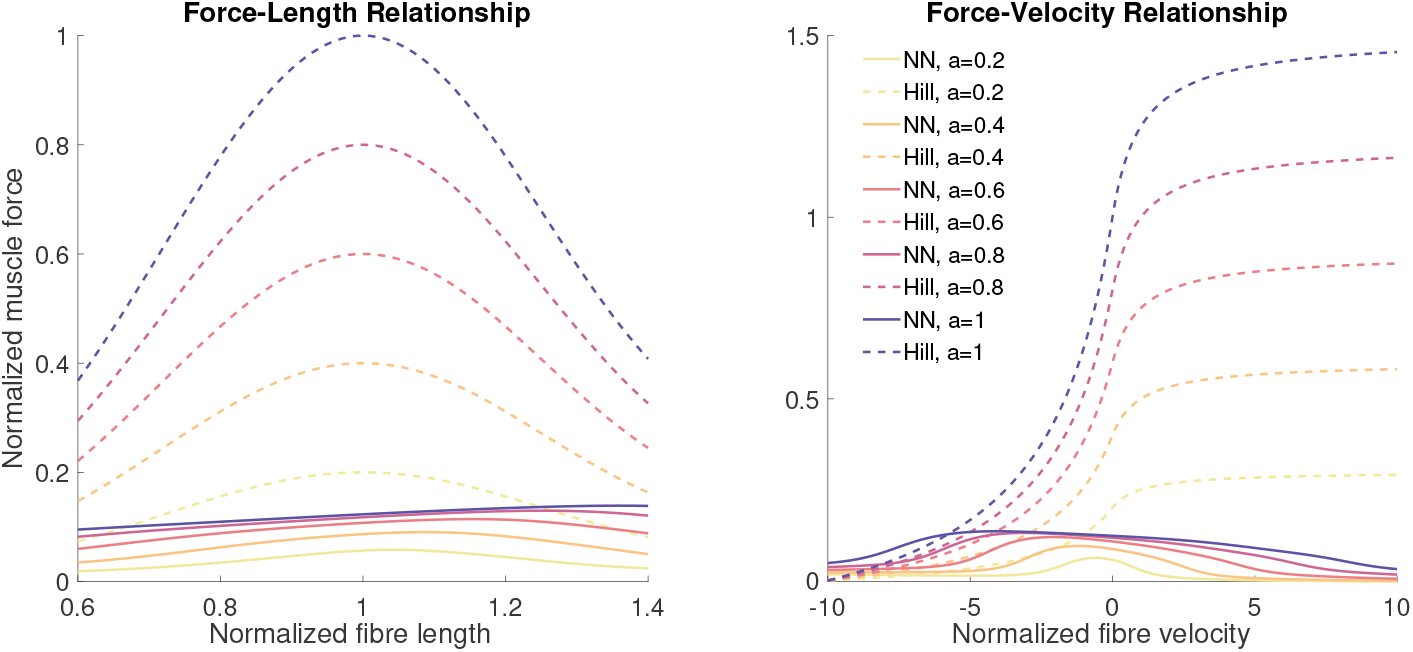
Force-length relationship, including parallel elastic element, and force-velocity relationship for NN-b1 and the Hill-type muscle model.

## 4 Discussion

We trained neural networks to estimate muscle force from activation, CE length, and CE velocity and investigated if these networks can estimate muscle forces more accurately than a Hill-type model, and how well they perform on unseen birds and muscles. We found that the neural networks generally outperformed Hill-type muscle models that were optimized using the same data as used for network training. The mean RMSE was lower for all test cases (Table 1), while mean coefficients of determination were higher (Table 2). In general, the validation and testing errors were of similar magnitude independent of dataset size used for training, but the smaller dataset led to more overfitting, as the difference between training and validation error was higher. We also found, both for the networks trained on smaller and larger datasets, that the neural network generally estimated force more accurately (lower mean RMSE and mean relative RMSE, and higher mean coefficient of determination) when applied to the same muscle as used for training, but on a different bird, compared to estimations on a different muscle on the same bird as used for training (Table 3, Fig. 3, Fig. 5, Supporting Fig. **??, ??**, and **??**). Furthermore, we found that our networks generally underestimated the maximum force amplification in the force-length and force-velocity relationships, but that they represented the force-length relationship at low activation levels well, while they represented the force-velocity relationship well only for shortening but not lengthening, which indicates that the networks did not perform well outside of the training data distribution (Fig. 6, Supporting Fig. **??, ??**, and **??**).

Our results show the potential of using neural networks to estimate muscle forces from muscle activation, muscle length, and muscle velocity. By training the neural networks on datasets that were recorded in dynamic conditions, we can ensure that the network better captures dynamic muscle behavior than Hill-type muscle models, which are based on limited, static conditions. These improved estimations are highlighted in Fig. 5, where the neural network estimates the smaller peaks that occur just before large peaks (e.g., at 10.3 and 11.6 s in (C)), which are generally not captured by Hill-type muscle models. These behaviors will be especially important in faster movements, such as running or change-of-direction movements, where the dynamic behavior of muscles is important (De Groote et al., 2016). However, to be able to implement a neural network, trained on guinea fowl data, to estimate muscle forces in a human musculoskeletal model, we need to have confidence that the network represents muscle mechanics correctly.

The reduced accuracy of the DF muscle predictions, as well as the underestimation of the maximum force amplification in the force-length and force-velocity relationships could be explained by inaccuracies in the maximum isometric force, in the activation scaling, or in the activation-deactivation time constants. Our dataset did not include tasks with maximal activation for normalization. Instead, we used the maximal activation over all (submaximal) trials, which might have led to a different normalization in the different muscles. This different normalization might have led to force underestimations and negative mean coefficients of determination, meaning that it is more accurate to use the mean than the force predictions, when the network was applied to another muscle (Fig. 3, Fig. 5, Supporting Fig. **??, ??**, and **??**), and a reduced maximum force amplification in the force-length and force-velocity relationships. The amplification factor was less than 0.3, instead of 1 for the force-length relationship and about 1.5 for the force-velocity relationship (Fig. 6). These low amplification factors could indicate that the activation scaling, to the maximum of the processed data, was not equivalent between the LG and the DF or that the maximum isometric force was not accurately estimated.

Our neural networks did not match the shape of the static force-length and force-velocity relationships either (Fig. 6, Supporting Fig. **??, ??**, and **??**). In the force-length relationship, the optimal fibre length increased with increasing activation, while previous experiments have shown that it decreases with increasing activation (e.g., Roszek and Huijing 1997; Rack and Westbury 1969; Schwaner et al. 2024). Nevertheless, a common observation between our most efficient network and those of Schwaner et al. (2024) is that at the highest activation, the force-length curve flattens out. Furthermore, there is a similar change observed between our network’s force-length relationship and that of Rack and Westbury (1969), where the ascending limb flattens out around optimal fibre length for higher activations. Furthermore, we found that the maximum shortening velocity decreased with decreasing activation, which was previously shown (Chow and Darling, 1999). However, the networks’ force-length-velocity relationships did not display the correct shape for the entire tested range. Only at low activation was the optimal fibre length displayed accurately, around 1 (optimal fibre length), while for larger activation, it was longer. Similarly, the force-velocity relationship did not display a force increase at lengthening velocities, though the shape for shortening velocities was relatively accurate. These results indicate that the networks’ output was inaccurate at the extremes of the training data, e.g., at large activation and large lengthening velocities (Fig. 6, Supporting Fig. **??, ??**, and **??**). This inaccuracy could be explained by a lack of training data in these regions, since most data was recorded at low and medium activation, and fibre lengths and velocities closer to optimal and isometric conditions, meaning that the networks did not perform well outside of the training data distribution. Another reason could be found in a recent study by Schwaner et al. (2024), who found that in-vivo force-length dynamics might not be represented well by in-situ measurements of the force-length relationship. Therefore, it is generally not unexpected that the force-length and force-velocity relationships did not match well, since our conditions did not meet the specific conditions in which these relationships apply. Yet, understanding what neural networks can learn about these relationships, based on in-vivo conditions, can uncover potentially valuable insights about muscle mechanics and performance outside the training distribution. However, we need to find additional approaches to further validate the neural networks for application across species.

To achieve better neural network performance, we recommend creating a more suitable training dataset. An ideal dataset, that is applicable to a broad range of activities and species, would include data of at least three different muscles and at least three different animal species, and includes different tasks that represent the full functional range of the animals as much as possible. By including three muscles, we can perform training on two, and test on the third muscle, which would avoid a bias in the network towards the single muscle used in training. Such a bias is likely present in our networks, which performed better on the muscle that they were trained on (LG) than the unseen muscle (DF). Furthermore, to apply such a network to a human musculoskeletal model, we need to at least validate the model on different species to test how well it performs on an unseen species. It would be best to directly validate on human data, but such a validation is impossible due to the need for invasive sensors. Again, by including at least three species, we can train on two and test on the third, and thereby avoid that the network biases towards a specific species. This ideal dataset should also include the full functional range of the animals as much as possible. To do so, both submaximal and maximal tasks should be included. These maximal tasks should lead to all measured muscles being maximally activated, such that the normalization is comparable between muscles. Furthermore, it should include recordings with a large distribution of muscle lengths, and shortening and lengthening velocities. This ideal dataset, to train a network as a replacement of the Hill-type muscle model, should not include data of fatigued muscles, since fatigue alters the relationship between activation and force over time. Eventually, fatigue could also be included in a neural network if a fatigue measurement can be included in the training data.

It is also important to realize that a neural network is only as good as the data that is used for its training. In practice, training datasets will always have limitations. For example, it could be that a neural network is well-suited to predict forces for walking when it is trained only on submaximal activation data, but not for jumping. Furthermore, there might be too much difference in data recordings between species and muscles to train a neural network that is applicable on all species and muscles from one large dataset. Therefore, a neural network might not be as flexible as a phenomenological model such as the Hill-type muscle model, though Hill-type muscle models are actually often applied even though they are known to be inaccurate (e.g., Abbott and Aubert 1952). However, the internal relationships of a Hill-type muscle model are understood, and it is therefore known that they still represent muscle mechanics to some extent. In contrast, we generally do not understand the internal behavior of a neural network, which acts like a black box, and therefore a neural network cannot be deployed without proper validation. At the same time, DeepLabCut, an effective machine learning model for pose estimation, has a similar black box behavior. This model can be used for different movements as well as different species through transfer learning with little additional training data (Nath et al., 2019). Such an approach could potentially also be used to transfer muscle models between species. However, transfer learning towards humans would again be challenging since direct measurements are impossible. Instead, it might be possible to create sufficient training data by estimating muscle forces through inverse kinematics and dynamics from optical motion capture, possibly combined with surface electromyography. In this approach, a Hill-type muscle model is normally used to estimate the muscle forces, which means that the neural network might in the end be trained to replicate a Hill-type muscle model instead of the actual muscle mechanics. Furthermore, the black box nature could be reduced through physics-informed training (e.g. Raissi et al. 2019; Zhang et al. 2022). In physics-informed training, the loss is calculated not only from training data, but also based on some physics relationships that the network should follow. For example, in case of a muscle model, a loss could be included based on how well the network follows the force-length and force-velocity relationships. Then, the neural network is not forced to follow those relationships exactly, but uses them to steer the training, while still also matching the training data. To achieve that, a delicate balance must be found between physics-based and data-based learning, such that physics models are used in conditions where they are known to apply, while the network learns from and adapts to the data in other conditions, potentially correcting for incorrect specifications in the physics model.

We also compared our results to other relevant muscle models (Whitney et al., 2022; Liu et al., 1999). Liu et al. (1999) successfully trained an artificial neural network to estimate muscle force in a cat soleus solely from electromyography signals (Liu et al., 1999). We compared our mean relative RMSE values to theirs in the most similar shared scenario: training on EMG and force data of the same muscle of several animals (cats or guinea fowl) and testing on another (cat or guinea fowl). Comparing the mean relative RMSE on the LG muscle of unseen birds for all four of our networks to their relative RMSE at speed 1.2 m s^-1^ (the highest speed of their dataset that is closest to the lowest speed of our dataset), we observe lower or comparable relative RMSE values (8.6%, 13.8% and 13.5% compared to 12% in Table 3 of Liu et al. 1999). Predictions on the LG of bird 5 using NN-b5 resulted in worse relative RMSE (29.6% compared to 12% in Table 3 of Liu et al. 1999). However, predictions on bird 5 are typically poor for all of our networks, regardless of the status of bird 5 as ‘seen’ or ‘unseen’ during training. When we used our data to train a model to estimate force from electromyography data only, the network performance was worse. This reduction could be caused by the fact that our dataset, which included perturbations, was specifically designed to include cases where a different force was created despite having the same muscle activation, while Liu et al. (1999) used steady-state walking data, where the relationship between muscle activation and force is more direct. Due to this training approach, their model will likely not generalize to non-steady conditions, as these are not included in their training data. We also compared our mean coefficient of determination values to a recently developed winding filament model (Whitney et al., 2022), which was optimized on the same data. Compared to this model, our mean coefficients of determination were comparable or higher for the LG data used in training (0.79, 0.67, and 0.80 compared to 0.81, 0.71, and 0.78 in Table 2 in Whitney et al. 2022), while they were much lower for the unseen bird (0.27 compared to 0.74 in Table 2 in Whitney et al. 2022). Also for the DF, the unseen muscle, our mean coefficients of determination were worse (0.49, 0.30, -0.08, and -0.13 compared to 0.82, 0.64, 0.64, and 0.48 in Whitney et al. 2022). However, Whitney et al. (2022) performed a model parameter optimization for each muscle and each trial individually, while we tested on unseen birds or muscles.

Since the dataset used here was not recorded specifically for neural network training, we could not test different approaches. For example, we suspect that the under- and overestimation of peak forces in the DF is related to a difference in activation scaling between the two muscles or inaccurate estimates of the maximum isometric force. We scaled the activation to the maximum activation recorded in the respective muscle over all trials of the same bird, but it is well possible that this is not equivalent between the muscles if less activation is required in one muscle than in the other for the recorded activities (e.g., 40% of the maximum for the LG and 80% of the maximum for the DF). Furthermore, the maximum isometric force was determined from the physiological cross-sectional area through the maximum stress, but the maximum stress might be different between muscles (Maganaris et al., 2001). Since the network is only trained on one muscle, it has likely developed a bias towards this muscle’s error in the activation and maximum isometric force, and therefore the mean RMSE and the mean relative RMSE are much lower for the LG than for the DF. Furthermore, this bias seems persistent among the same muscle of different birds, but not between different muscles of the same bird. When training on two or more muscles, such a bias could be avoided. However, since data of only two muscles was available, we could not have tested this scenario on independent data.

In this work, we trained neural networks to be similar to a Hill-type muscle model, but in the future, different network architectures could be explored to further improve network performance and have them better represent muscle mechanics. For example, there is a history dependence in muscle force generation (Herzog, 2004) which could be captured in a recurrent neural network. Furthermore, neural networks could also be used as a tool for data-based muscle mechanics exploration. By comparing the quality of different architectures or input combinations, the relevance of different muscle states or parameters to muscle force generation could be explored to formulate hypotheses about muscle mechanics. These hypotheses could then be further tested in specific experiments.

In conclusion, we found that a neural network can estimate muscle force more accurately than a Hill-type muscle model, both on different muscles and birds, and independent of training dataset size. However, a larger dataset seemed to reduce overfitting towards the training data. We also found that, when training on data of one muscle, neural networks make better estimations on the same muscle of another bird, than on another muscle of the same or a different bird. Future studies should further investigate if this effect can be mitigated by training on more than one muscle, which reduces the bias towards the muscle used for training, while we have also made recommendations on an ideal dataset including maximum activation tasks which could also mitigate this issue. In conclusion, we have shown that neural networks are a promising approach to developing new muscle models, and they could also be a helpful tool when creating new hypotheses about muscle mechanics.

## List of Symbols and Abbreviations

A_*rel*_: Curvature of the force-velocity relationship
c_3_: Continuity factor ensuring a smooth transition in the force-velocity relationship
CE: Contractile element
DF: Digital flexor
F_*CE*_: Contractile element force
F_*Hill*_: Tendon or muscle force calculated by Hill-type model f(l_*CE*_) Force-length relationship
F_*meas*_: Experimentally measured tendon or muscle force F_*P EE*_ Parallel elastic element force
g_*max*_: Maximum force amplification during muscle lengthening
g(v_*CE*_): Force-velocity relationship
Hill-r01: Hill-type muscle model optimized on the first trial of bird 1, at 1.8 m/s and a level surface
Hill-r12: Hill-type muscle model optimized on the twelfth trial of bird 1, at 4.5 m/s with 7 cm obstacles
k_*PEE*_: Stiffness of the parallel elastic element
l_*CE*_: Muscle or contractile element length
LG: Lateral gastrocnemius
l_*slack,P EE*_: Slack length of the parallel elastic element
NN-b1: Neural network trained on a large dataset with trials of varying obstacle heights (level, 5 and 7 cm) and speeds (1.8, 3.0, 3.5, 3.8 and 4.5 m/s). Training excluded all trials of bird 1
NN-b2: Neural network trained on a large dataset with trials of varying obstacle heights (level, 5 and 7 cm) and speeds (1.8, 3.0, 3.5, 3.8 and 4.5 m/s). Training excluded all trials of bird 2
NN-b3: Neural network trained on a large dataset with trials of varying obstacle heights (level, 5 and 7 cm) and speeds (1.8, 3.8 and 4.5 m/s). Training excluded all trials of bird 3
NN-b5: Neural network trained on a large dataset with trials of varying obstacle heights (level, 5 and 7 cm) and speeds (1.8, 3.0, 3.5, 3.8 and 4.5 m/s). Training excluded all trials of bird 5
NN-r01: Neural network trained on the first trial of bird 1, at 1.8 m/s and a level surface
NN-r12: Neural network trained on the twelfth trial of bird 1, at 4.5 m/s with 7 cm obstacles PEE Parallel elastic element
R^2^: Coefficient of determination
RMSE: Root-mean-square error RMSE% Relative root-mean-square error SEE Series elastic element
v_*CE*_: Muscle or contractile element velocity
v_*CE,max*_: Maximum shortening velocity of the force-velocity relationship W Width of the force-length relationship

## Author contributions

MAD and ADK conceptualized the study, MEA and MAD performed data curation, MEA and ADK performed formal analysis, ADK acquired funding, MEA and ADK performed the investigation and developed the methodology, ADK administered the project, MAD provided resources, MEA developed the software, ADK supervised the study, MEA and ADK validated the study and visualized the results. MEA and ADK wrote the original draft, and MEA, MAD, and ADK reviewed and edited the draft.

## Data availability

All relevant code and data are available on Zenodo: [https://doi.org/10.5281/zenodo.14512831] and [https://doi.org/10.5281/zenodo.14727454].

