## Supporting Document for "Neural Networks Estimate Muscle Force in Dynamic Conditions Better than Hill-type Muscle Models"

### Results

This supplemental document shows Fig. 4 and Fig. 5 for the networks NN-b2, NN-b3, and NN-b5.

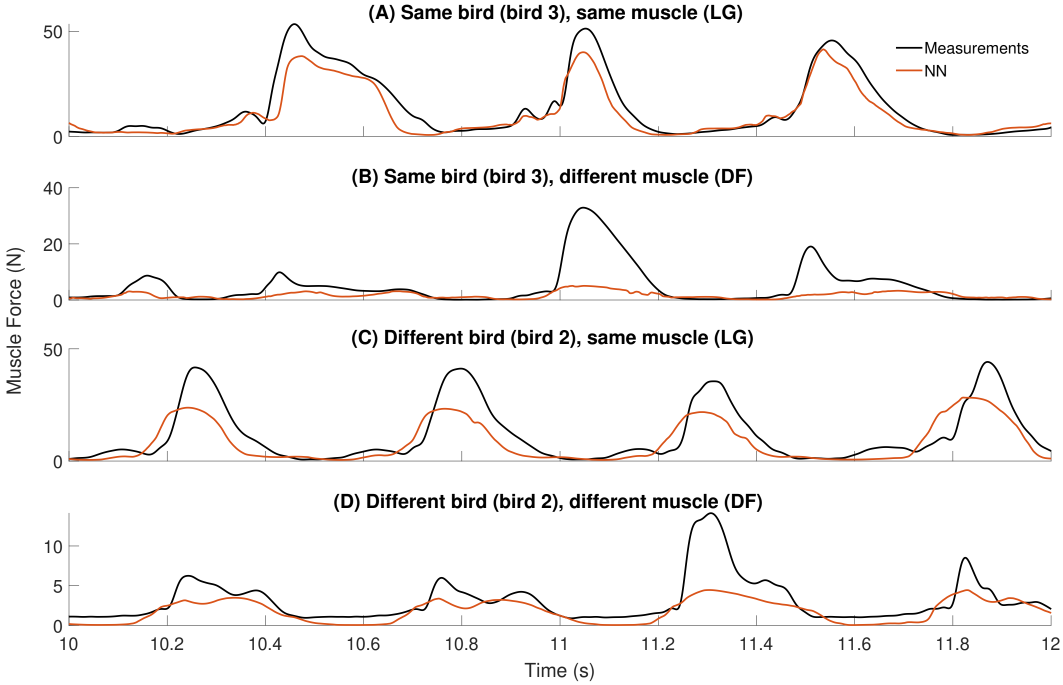


Figure S1: Muscle force predictions by NN-b2 and measurements over time, for part of a trial with speed 1.8 m $s^{-1}$ and 7 cm elevation for (A) the same bird and muscle as used for network training, (B) the same bird but a different muscle, (C) the same muscle of a different bird, and (D) a different muscle of a different bird.

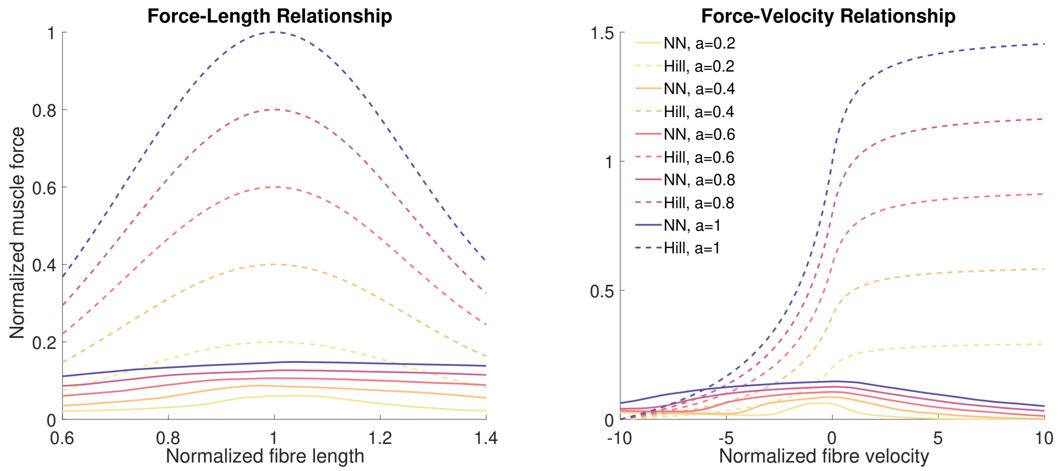


Figure S2: Force-length relationship, including parallel elastic element, and force-velocity relationship for NN-b2 and the Hill-type muscle model.

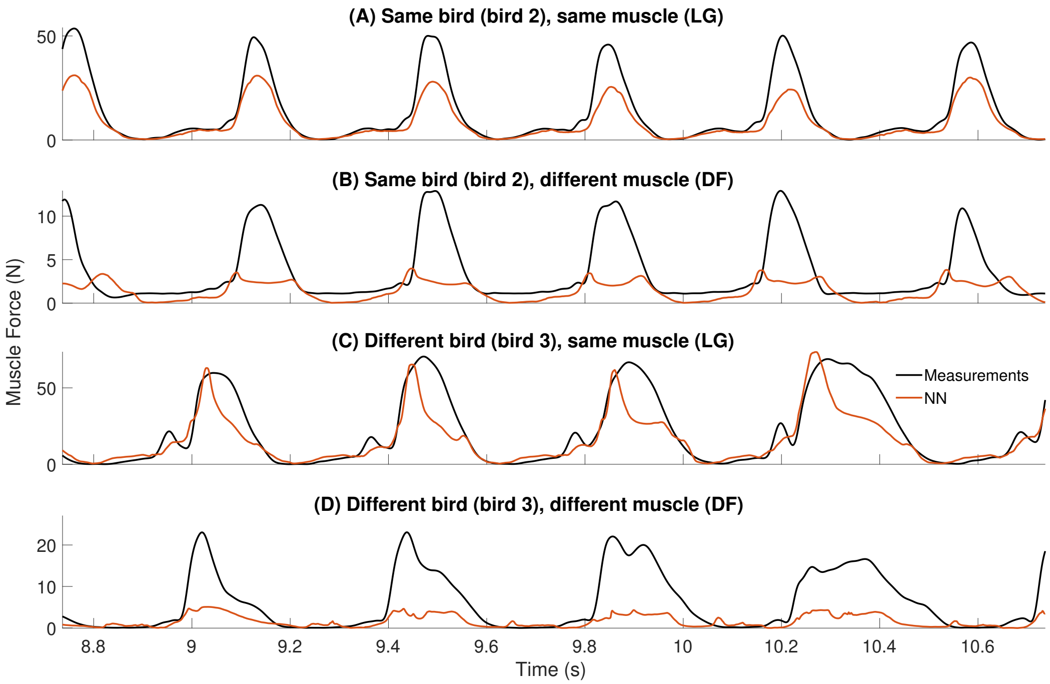


Figure S3: Muscle force predictions by NN-b3 and measurements over time, for part of a trial for (A) the same bird and muscle as used for network training, (B) the same bird but a different muscle, (C) the same muscle of a different bird, and (D) a different muscle of a different bird. The presented trials do not include an obstacle and their speeds are 3.8 and 3.5 m $s^{-1}$, for bird 2 and 3, respectively. Despite their speed difference, these trials allowed for the closest comparison of test scenarios, among the reserved test trials.

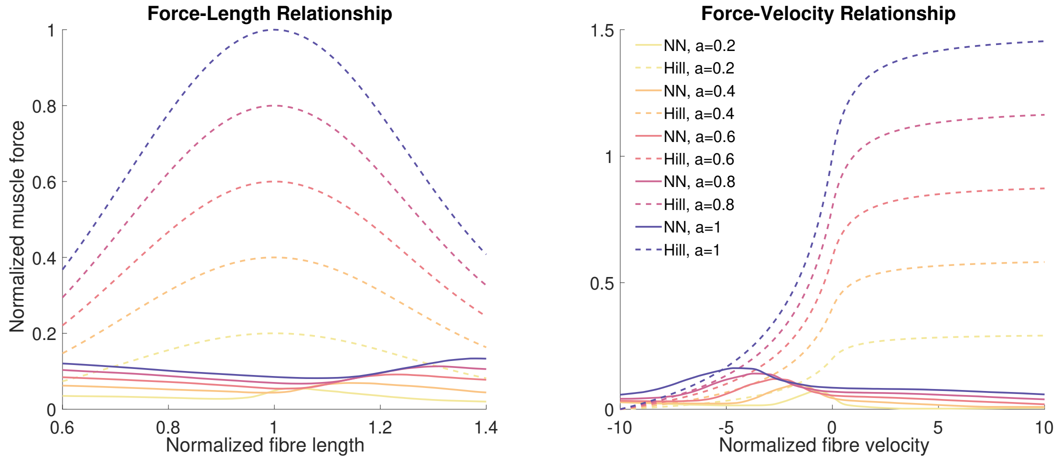


Figure S4: Force-length relationship, including parallel elastic element, and force-velocity relationship for NN-b3 and the Hill-type muscle model.

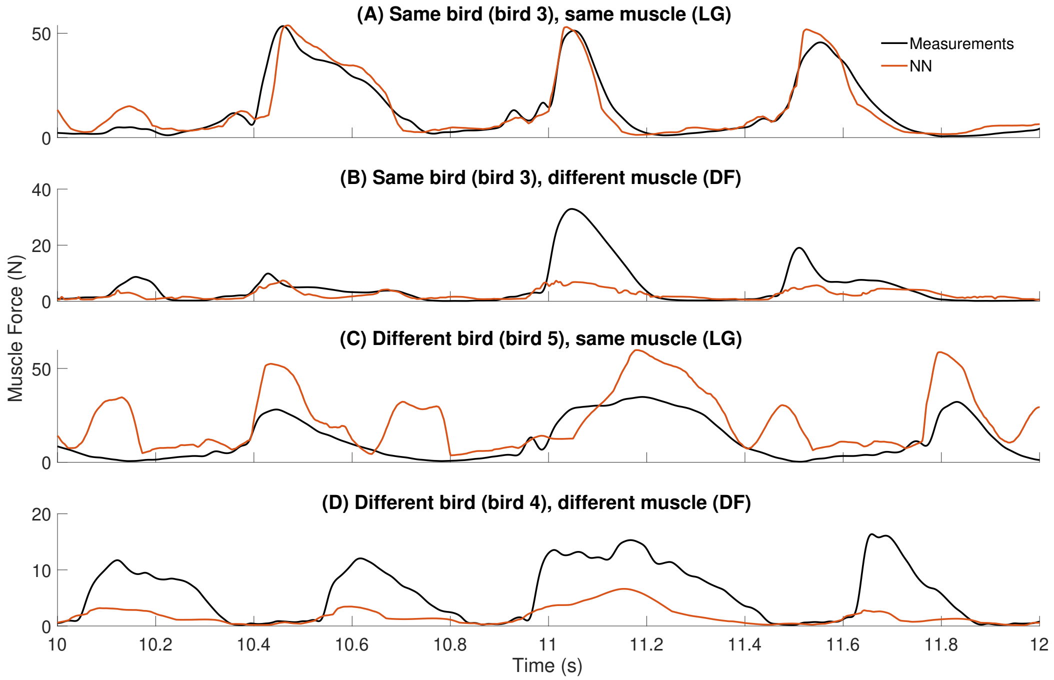


Figure S5: Muscle force predictions by NN-b5 and measurements over time, for part of a trial with speed 1.8 m $s^{-1}$ and 7 cm elevation for (A) the same bird and muscle as used for network training, (B) the same bird but a different muscle, (C) the same muscle of a different bird, and (D) a different muscle of a different bird.

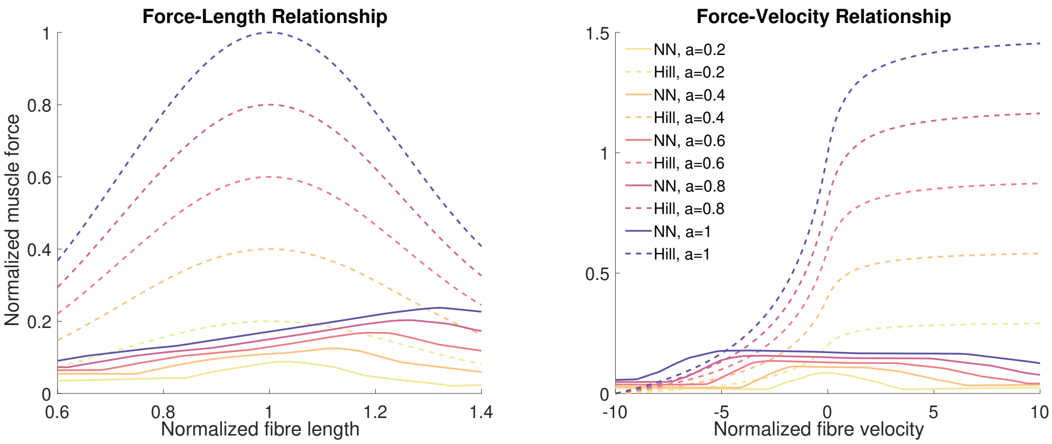


Figure S6: Force-length relationship, including parallel elastic element, and force-velocity relationship for NN-b5 and the Hill-type muscle model.
